# Bacteria-mediated stabilization of murine norovirus

**DOI:** 10.1101/2022.01.21.477311

**Authors:** Melissa R. Budicini, Julie K. Pfeiffer

**Affiliations:** Department of Microbiology, University of Texas Southwestern Medical Center, Dallas, Texas, USA

**Keywords:** Murine norovirus, stability, bacteria

## Abstract

Enteric viruses encounter various bacteria in the host which can impact infection outcomes. The interactions between noroviruses and enteric bacteria are not well understood. Previous work determined that murine norovirus (MNV), a model norovirus, had decreased replication in antibiotic-treated mice compared with conventional mice. Although this suggests that microbiota promote MNV infection, the mechanisms are not completely understood. Additionally, prior work with other enteric viruses such as poliovirus and coxsackievirus B3 demonstrated that virions bind bacteria, and exposure to bacteria stabilizes viral particles and limits premature RNA release. Therefore, we examined interactions between MNV and specific bacteria and the consequences of these interactions. We found that the majority of Gram-positive bacteria tested stabilized MNV, while Gram-negative bacteria did not stabilize MNV. Both Gram-positive and Gram-negative bacteria bound to MNV. However, bacterial binding alone was not sufficient for virion stabilization since Gram-negative bacteria bound MNV but did not stabilize virions. Additionally, we found that bacterial conditioned media also stabilized MNV and this stabilization may be due to a small heat stable molecule. Overall this work identifies specific bacteria and bacterial components that stabilize MNV and may impact virion stability in the environment.

**Importance:** Enteric viruses are exposed to a wide variety of bacteria in the intestine, but effects of bacteria on viral particles are incompletely understood. We found that murine norovirus (MNV) virion stability is enhanced in the presence of several Gram-positive bacterial strains. Virion stabilizing activity was also present in bacterial culture medium, and activity was retained upon heat or protease treatment. These results suggest that certain bacteria and bacterial products may promote MNV stability in the environment, which could influence viral transmission.

## Introduction

Noroviruses are a leading cause of nonbacterial gastroenteritis disease around the world. Human norovirus (HuNoV) causes 1 billion infections and 200,000 deaths annually (1–3). The economic impacts of these infections are significant, with $4.2 billion of direct health system costs and $60.3 billion in societal costs globally each year (1, 4). Despite its impact, little is known about the mechanisms of disease of HuNoV due to the lack of a robust cell culture system or small animal model. However, in 2003 a genetically related virus, murine norovirus (MNV), was discovered and is now used as a model system for HuNoV due to its genetic similarity, efficient replication in vitro, and tractable mouse models (5, 6). MNV is a small non-enveloped, positive-sense RNA virus with a 7.5-kb genome (7). The main site of MNV infection is in the intestine and it is spread through the fecal oral route.

During infection in the intestine, enteric viruses encounter 10^14^ bacteria (8). Enteric viruses such as poliovirus, coxsackievirus, and reovirus can interact directly with the gut microbiota to enhance infection through a variety of mechanisms such as increased host cell binding, increased receptor binding, and increased viral stability in the presence of bacteria (9–15). For MNV, depletion of microbiota by antibiotic treatment in mice decreases viral titers during acute and persistent infection (16, 17). Certain host genes involved in innate immune responses, including the interferon lambda receptor, Stat1, and Irf3, were required for antibiotic-mediated loss of viral persistence in mice, suggesting that microbiota promote MNV persistence by inhibiting host innate responses (17). However, effects of microbiota on MNV are complex and site-specific, since microbiota can inhibit MNV infection of the upper intestine via bile acid priming of interferon lambda responses (18). MNV can bind directly to bacteria in vitro (19, 20). Although these finding suggest that the presence of bacteria is important for promoting MNV infection, the mechanisms are incompletely understood.

Because MNV is spread through the fecal oral route, virion stability in the environment is necessary for maintenance of viral infectivity and transmission to a new host. Viral stability can be measured by exposing viral particles to high temperatures and quantifying remaining viable viruses. Heat causes changes in viral capsid conformations that can lead to viral genome release and virion inactivation (21). For poliovirus, another enteric virus spread through the fecal oral route, the presence of bacteria and bacterial components such as lipopolysaccharide (LPS) can stabilize the virus capsid which leads to increased transmission (10). In this study we determined the effect of bacteria and bacterial components on MNV thermostability. We used heat inactivation assays with whole bacteria, bacterial surface molecules, and bacterial conditioned media and determined their impact on viral stability. We found that specific Gram-positive bacteria and conditioned media from Gram-positive bacteria stabilized MNV. Conversely, the Gram-negative bacteria tested had little impact on viral stability. Overall, these findings define interactions between MNV and specific bacteria which may provide insight into virion environmental stability and transmission.

## Results

### Most Gram-positive bacteria tested enhance stability of MNV

To determine if bacteria can stabilize MNV, we exposed virions to different bacterial strains and quantified viral infectivity following heat exposure. MNV and other non-enveloped RNA viral particles can be inactivated at high temperatures due to premature genome release. Previous studies have shown that bacteria and bacterial compounds are able to stabilize other non-enveloped RNA viruses at high temperatures (10–12). We first tested viral stability following incubation at 42°C for 6 hours, a condition which inactivates approximately 90% of MNV infectivity (Fig. 1). The virus was mixed with PBS, streptavidin beads, or bacteria and incubated at 42°C for 6 hours. Plaque assays were used to determine the amount of viable virus remaining compared to samples incubated at 4°C. The majority of Grampositive bacterial strains increased the amount of viable virus compared with PBS or bead control (Fig. 1). However, all of the Gram-negative bacteria tested had no significant impact on viral stability as compared with PBS (Fig. 1). We next examined stabilization of MNV at a higher temperature. The virus was mixed with PBS, *S. aureus,* or *E. saccharolyticus* at 46°C for 4 hours. The stabilization effect of Gram-positive bacteria was also evident at this higher temperature (Fig. 2A). The 4 hour incubation at 46°C was used for all subsequent experiments due to the increased dynamic range provided by these conditions.

**Figure 1.**
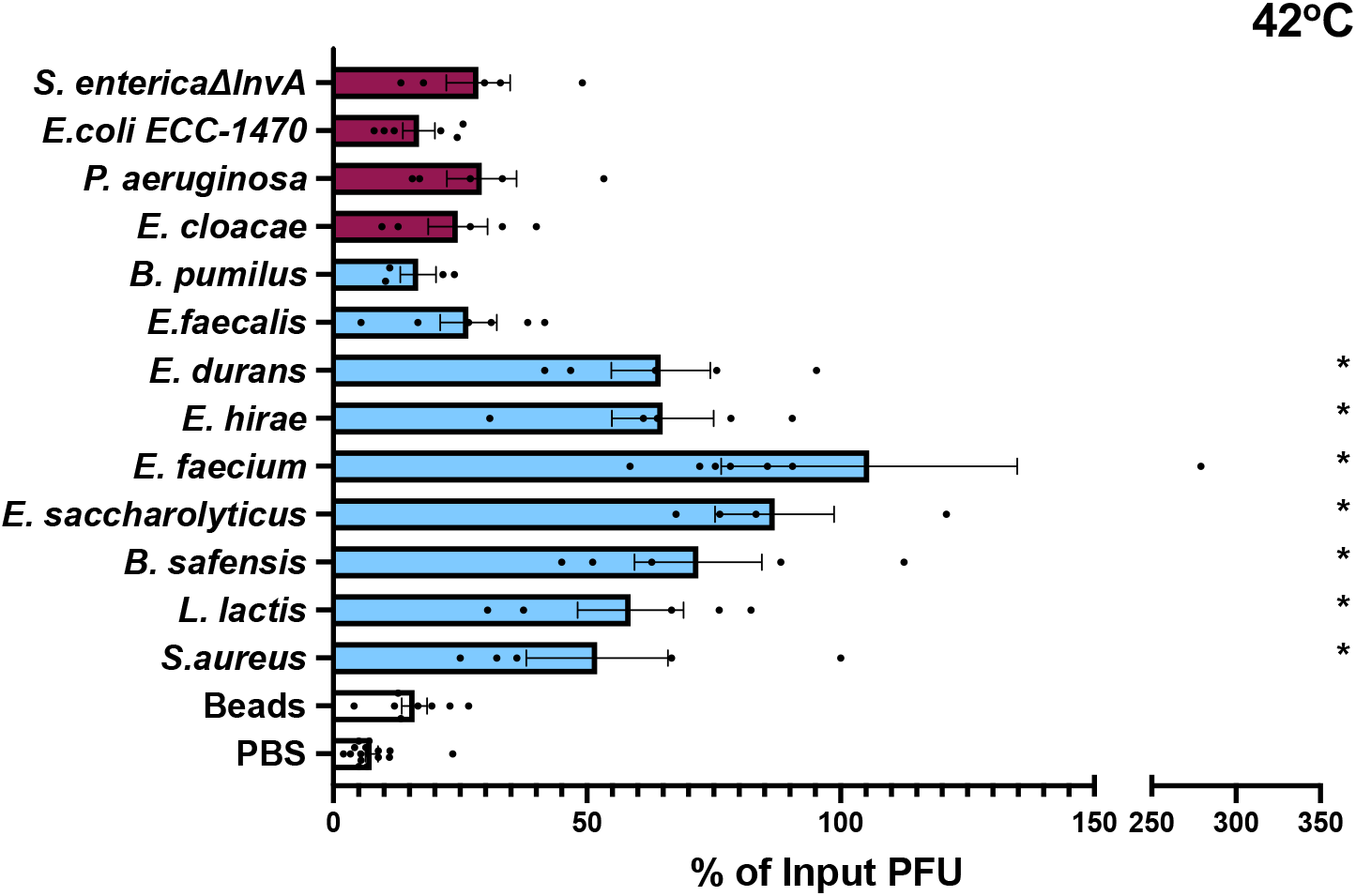
Effects of bacteria on MNV stability. Thermal stability assays were performed by incubating 1 x 10^6^ PFU MNV with PBS, strepdavidin beads, or 1 x 10^9^ CFU bacteria at 42°C for 6 h. The amount of viable virus after each assay was determined by plaque assay and compared to a 4°C PBS viral titer to calculate percent of input PFU that remained. Data points are averaged two replicates per experiment from 4 to 16 independent experiments *(n* = 4 to 16). Bars are shown for SEM. Statistical significance was determined by one-way ANOVA compared with the PBS samples (*, *P* < 0.05). Clear bars= controls, blue bars= Gram-positive bacteria, purple bars= Gram-negative bacteria.

We also determined whether bacterial surface molecules could stabilize MNV. We included surface molecules from both Gram-positive and Gram-negative bacteria as well as some non-bacterial glycans that have previously been shown to stabilize other enteric viruses (10). We incubated MNV with either PBS or 1 mg/mL of each molecule at 46°C for 4 hours. We found that lipoteichoic acid (LTA), a surface molecule from Gram-positive bacteria, was able to stabilize MNV. Interestingly we found that lipopolysaccharide (LPS), a Gram-negative surface molecule, was also able to stabilize MNV (Fig. 2B). Overall these data suggest that LTA may contribute to MNV stabilization by Gram-positive bacteria, but that other molecules may also be sufficient for stabilization.

**Figure 2.**
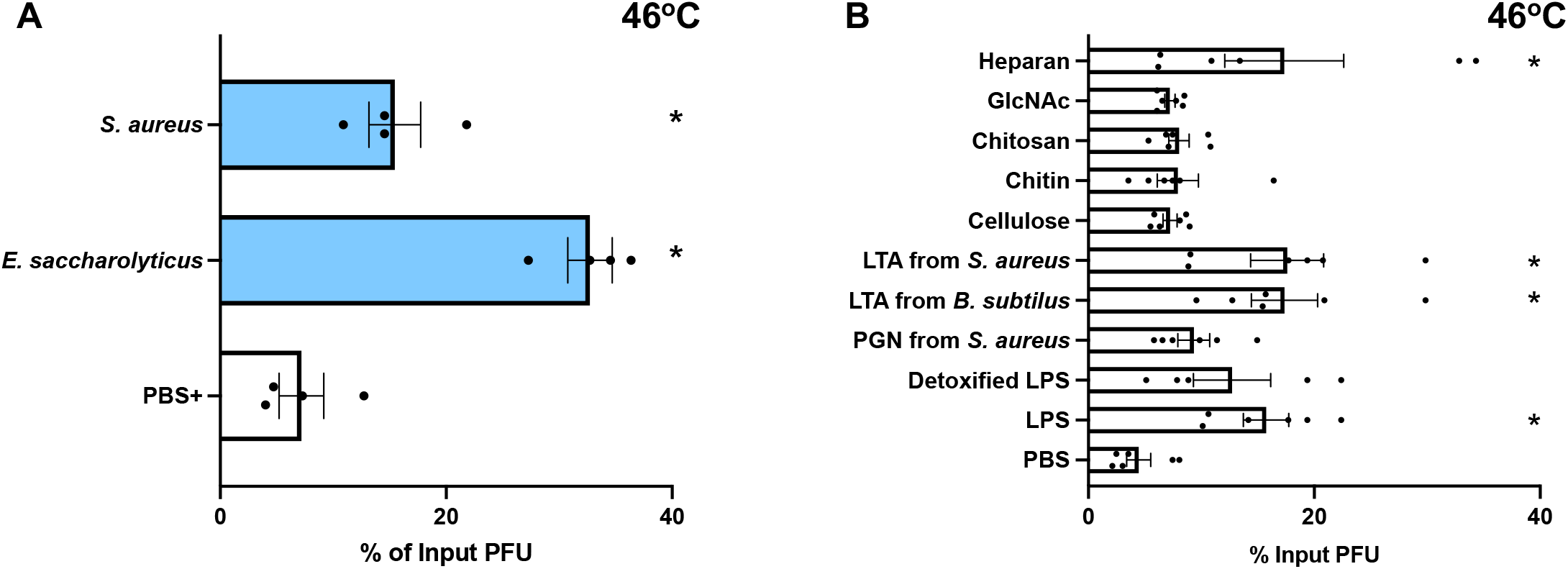
Effects of bacteria and compounds on MNV stability at high temperature. Thermal stability assays were performed by incubating 1 x 10^6^ PFU MNV with PBS, strepdavidin beads, or 1 x 10^9^ CFU bacteria, or 1mg/mL compounds at elevated temperatures. The amount of viable virus after each assay was determined by plaque assay and compared to a 4°C PBS viral titer to calculate percent of input PFU that remained. (A) Viral incubation with bacteria at 46°C for 4 h. Data are representative of 3 independent experiments (*n =* 6). (B) Viral incubation with compounds at 46°C for 4 h. GlcNAc, N-acetylglucosamine; LPS, lipopolysaccharide; LTA, lipoteichoic acid; PGN, peptidoglycan. Data are representative of 3 independent experiments (*n*= 5 to 6). Bars are show mean and SEM. Statistical significance was determined by one-way ANOVA compared with the PBS samples (*, *P* < 0.05).

### MNV binds to both Gram-positive and Gram-negative bacteria

Since bacteria were able to stabilize MNV, we determined whether MNV could interact directly with bacteria. Previously it was shown that MNV can bind to certain bacteria, although the consequences of these interactions was unclear (19). One hypothesis for the increased stabilization effects of Grampositive bacteria is that these bacteria simply bind MNV more efficiently. We used bacterial binding assays to determine whether the increase in stabilization by Gram-positive bacteria was a result of increased binding as compared to Gram-negative bacteria. To quantify binding, ^35^S-labeled MNV was incubated for 1 hour with either beads or a subset of the bacteria, followed by centrifugation, washing, and scintillation counting of the bacterial pellets to determine the percent of virus bound to the bacteria. We found that MNV binds to both Gram-positive and Gram-negative bacterial strains (Fig. 3). This indicates that not all viral binding to bacteria leads to stabilization. For example, *P. aeruginosa* bound to MNV but failed to stabilize in thermostability assays. Additionally, the increased viral stabilization by Gram-positive bacteria is not a consequence of higher binding. Overall, these results indicate that MNV can bind to both Gram-positive and Gram-negative bacteria, and although binding may be required for stabilization it is not sufficient for stabilization.

**Figure 3.**
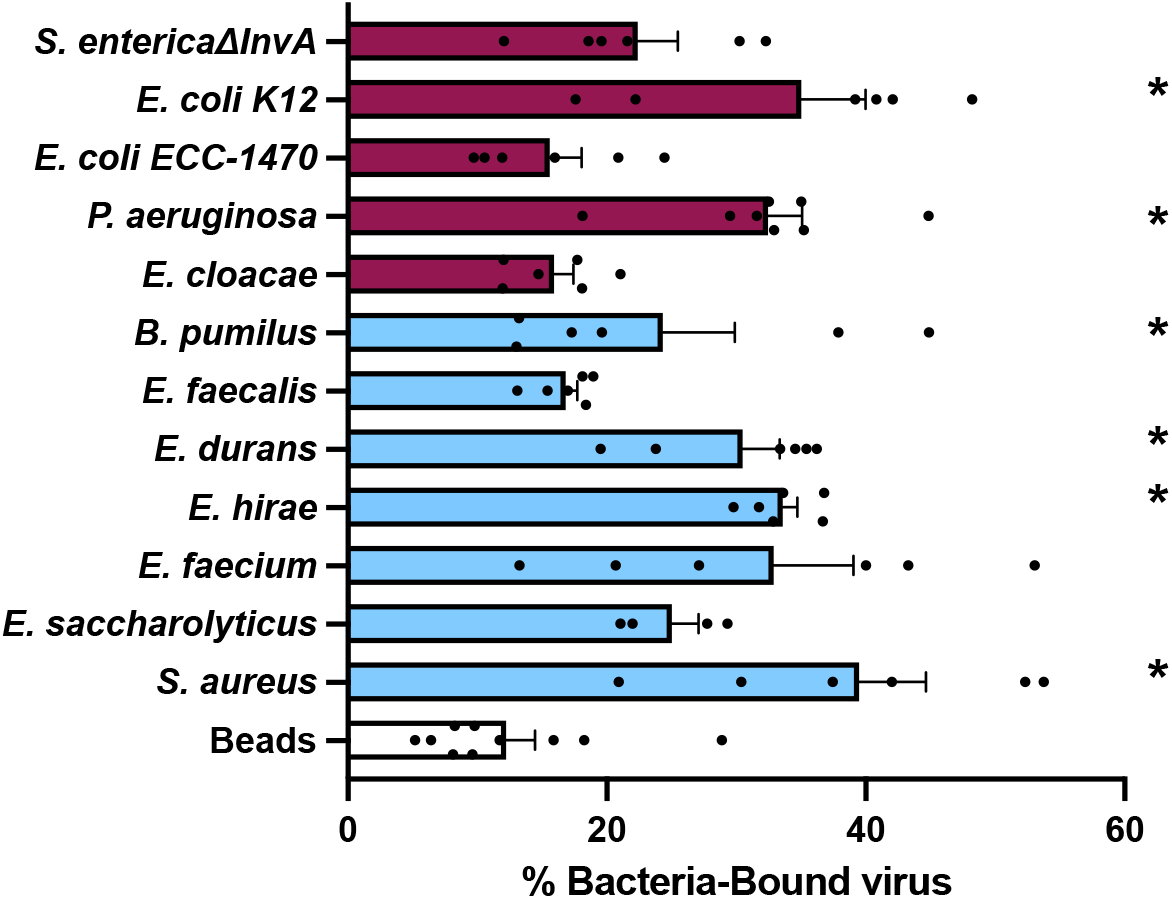
MNV binds to bacteria. ^35^S-labeled viruses were incubated with 1 x 10^9^ CFU of bacteria for 1 h at 37°C. After incubation, bacteria were spun down and washed to remove free virus. Bound virus was quantified by scintillation counting. Data are representative of 2 to 4 independent experiments (*n* = 4 to 8). Bars show mean and SEM. Statistical significance was determined by one-way ANOVA compared with beads (*, *P* < 0.05). Clear bars= control, blue bars= Gram-positive bacteria, purple bars= Gram-negative bacteria.

### Incubation with bacteria does not impact MNV infectivity

We wanted to determine whether bacteria, in addition to stabilizing viral particles, enhance infectivity of virions in the absence of excess heat treatment. MNV was incubated with either PBS, *E. saccharolyticus, or S. aureus* at 37°C for 1 hour before infecting cells for a plaque assay. For this experiment the virus was incubated on the monolayer for 1, 5, or 15 minutes instead of the traditional 30 minute incubation time to determine if there were any infectivity differences in more stringent conditions. We found that there were no significant differences in titer for viruses incubated with PBS or bacteria at any time point (Fig. 4). This may indicate that bacteria can prevent viral particles from becoming inactivated at high temperatures, but may not alter the particles in a way that increases their ability to infect cells. Overall, these results indicate that incubation with bacteria does not make MNV more infectious for BV2 cells.

**Figure 4.**
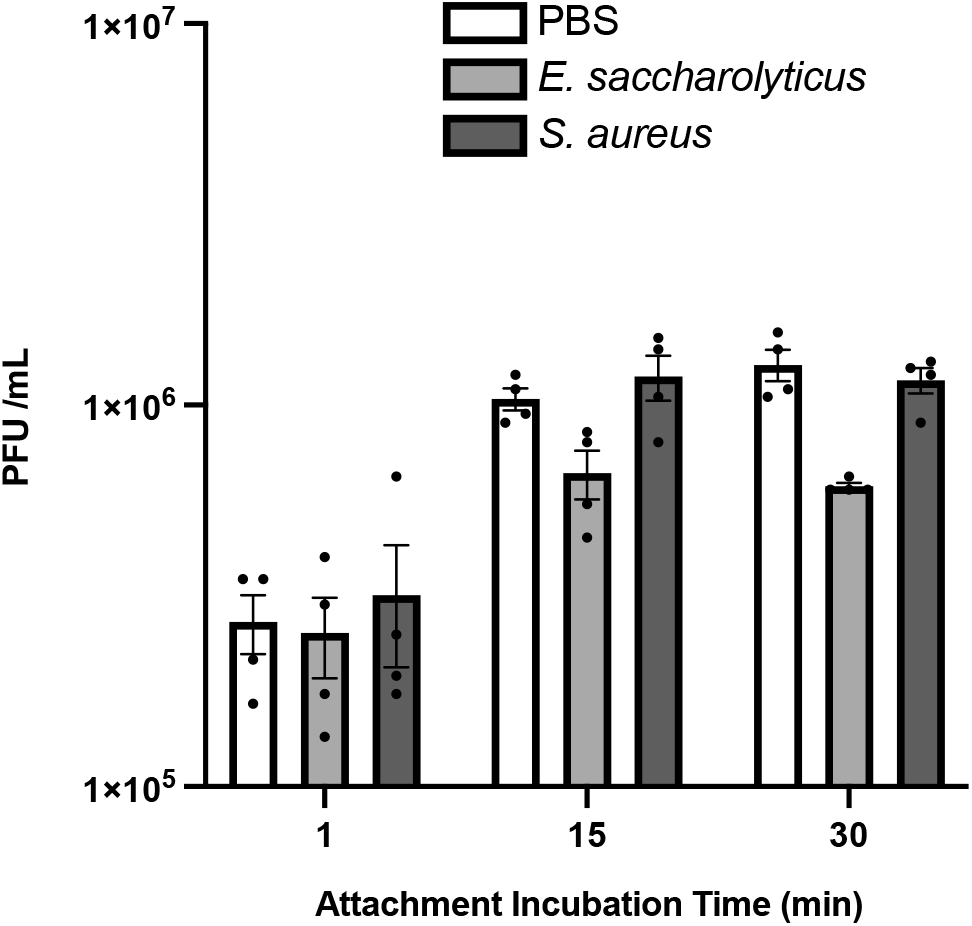
Effects of bacteria on MNV infectivity. Infectivity assays were performed by incubating 1 x 10^6^ PFU MNV with PBS or 1 x 10^9^ CFU bacteria at 37°C for 1 h prior to performing plaque assay. Each sample was either incubated for 1, 15, or 30 minutes on the cell monolayer during the attachment period. The amount of plaques was compared to the PBS condition for each timepoint. Data are representative of 2 independent experiments (*n = 4).* Bars show mean and SEM. Statistical significance was determined by two-way ANOVA compared to PBS (*, *P* < 0.05).

### Bacterial conditioned media from Gram-positive bacterial cultures can stabilize MNV

After observing that Gram-positive bacteria and bacterial surface molecules were able to stabilize MNV, we determined whether bacterial conditioned media from these strains also stabilized MNV. Bacterial conditioned media contains secreted factors or surface molecules that have sloughed off and these components could contribute to stabilization. Bacterial conditioned media can have a variety of effects on mammalian cells and eukaryotic organisms (22–24). However, the impact of conditioned media on viruses is unknown. We first tested viral stability in the presence of conditioned media for two strains, *E. saccharolyticus* and *S. aureus,* that had significant levels of stabilization at 46°C from whole bacteria (Fig. 2A). We found that spent media that was filtered with a 0.2 micron filter and boiled for 30 minutes was able to stabilize MNV when compared with BHI growth media alone, indicating that the stabilizing factor is heat stable and smaller than a whole bacterium (Fig. 5A). We then tested viral stability in the presence of conditioned media from a larger subset of bacteria. We found that the conditioned media from some Gram-positive bacteria was able to stabilize MNV. The conditioned media from all of the Gram-negative bacteria tested did not have an effect on viral stability, consistent with the data from the whole bacteria stability assay (Fig. 5B). Overall, these results indicate that the presence of whole bacteria is not required for stabilization and the stabilizing component is heat stable.

### A small, protease- and heat-stable molecule from bacterial conditioned media is sufficient to stabilize MNV

Since conditioned media from most Gram-positive bacterial strains stabilized MNV, we wanted to further define properties of the stabilizing factor. We chose the bacterial strains *E. saccharolyticus* and *S. aureus* from the Gram-positive group as two representative bacteria. Recent evidence shows that Gram-positive bacteria can produce extracellular vesicles, lipid bilayer enclosed particles that can contain diverse cargo such as nucleic acids, effector proteins, enzymes (25, 26). In order to determine if extracellular vesicles or other relatively large structures impacted MNV stability, we performed ultracentrifugation and used both the pellet and supernatant in thermal stability assays. We found that, for both strains, the supernatant but not the pellet was able to stabilize MNV, indicating that the stabilizing factor may not involve extracellular vesicles or other larger structures (Fig. 6). Next, we used size exclusion spin columns to fractionate the conditioned media. We tested both the >50kD and <50kD fractions in a thermal stability assay and found that for both strains there was significant MNV stabilization from the <50kD fraction. However, for *S. aureus* there was also statistically significant stabilization from >50kD fraction (Fig. 6). This may suggest that the stabilizing factor in the conditioned media can be a variety of sizes. Lastly, we determined whether the stabilizing factor in conditioned media is protease sensitive. Bacterial conditioned media was treated with proteinase K for 18 hours before the thermal stability assay with MNV. We found that for both strains the conditioned media maintained the ability to stabilize MNV after protease treatment. This suggests that the stabilizing molecule in the conditioned media is a heat- and protease-stable molecule that is relatively small.

**Figure 5.**
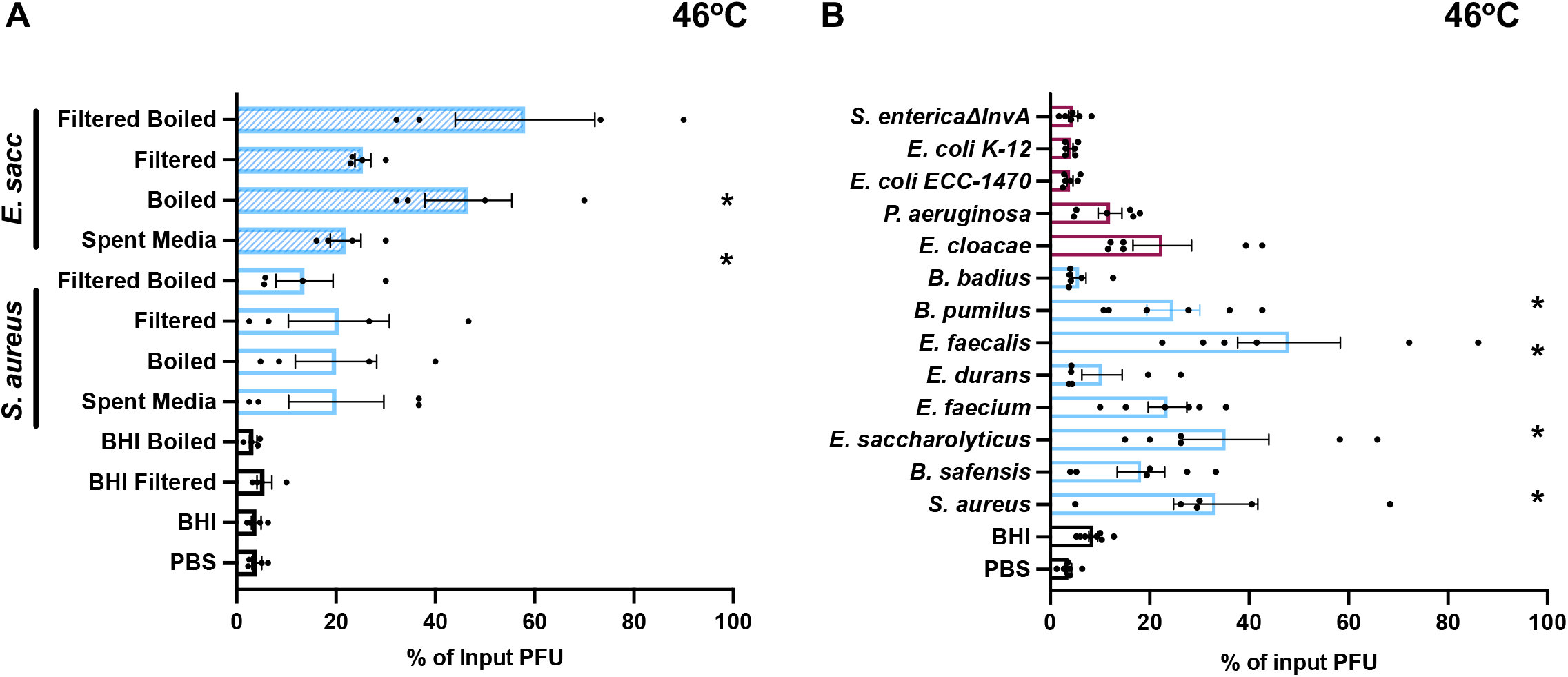
Effects of bacterial conditioned media on MNV stability. Thermal stability assays (46°C for 4 h) were performed by incubating 1 x 10^6^ PFU MNV with PBS, BHI media, or spent bacterial culture medium subjected to several different processing steps. (A) The thermal stability of MNV in spent media from *S. aureus* and *E. saccharolyticus* either boiled for 30 minutes at 95°C or passed through a 0.2 μm filter. (B) The thermal stability of MNV in conditioned media (filtered spent media) from different bacterial strains. For both A and B, the amount of viable virus remaining was compared to a 4°C PBS control to calculate percent of input PFU that remained. Data are representative of 3 to 4 independent experiments (*n* = 6 to 8). Bars show mean and SEM. Statistical significance was determined by one-way ANOVA compared to PBS (*, *P* < 0.05).

## Discussion

While it is established that the gut microbiota can influence MNV infection *in vivo* (16–18) there are still outstanding questions regarding the role of specific bacteria and underlying mechanisms. Previous studies have shown that pro-viral effects of the gut microbiota on MNV infection is lost in mice lacking certain innate immune factors. This suggests that the gut microbes may be altering the immune system in a way that promotes viral infection (17). In addition to the role that bacteria may play in modulating the immune response, we are interested in the direct effects of bacteria on MNV infection. Here we show that certain bacterial species, bacterial surface molecules, and bacterial conditioned media can increase the stability of MNV.

Our data indicate that Gram-positive but not Gram-negative bacteria are able to stabilize MNV. We found that most Gram-positive bacterial strains were able to stabilize MNV (Fig. 1). Although the mechanism underlying this stabilization is unclear, we found that LTA isolated from Gram-positive bacteria (*S. aureus* and *B. subtilus)* was able to stabilize MNV on its own (Fig. 2B). Since LTA is an important cell wall polymer in Gram-positive bacteria, these data suggest that it may play a role in stabilization (27).

Although we found that most Gram-positive bacterial strains stabilized MNV while Gram-negative did not (Fig. 1), we found that both groups were able to bind MNV (Fig. 3). We hypothesize that binding is a minimum requirement for stabilization, as all of the strains that were able to stabilize MNV also bound to the virus. However, we hypothesize that MNV binding is not sufficient for stabilization, as illustrated by the Gram-negative bacterial strains. Work with other enteric viruses in the picornavirus family also supports these ideas. Erickson et al. (11) and Dhalech et al. (13) showed that poliovirus or coxsackievirus B3 binding to bacteria does not always correlate with the ability of the bacteria to increase infectivity or stability. In addition to whole bacteria stabilizing MNV by a potential direct interaction mechanism, we have also shown that bacterial conditioned media is also able to stabilize MNV (Fig. 5). We found that the bacterial conditioned media from most Gram-positive bacteria was able to stabilize MNV (Fig. 5). Further, we found that for the conditioned media from *E. saccharolyticus* and *S. aureus,* the highest amount of stabilizing activity was found in the fraction of media with molecules <50kD and the stabilizing effect was maintained after protease treatment, suggesting the relevant factor is not a protease-sensitive protein (Fig. 6). Further work is needed to determine the exact identity and biochemical properties of the stabilizing components of bacterial conditioned media.

**Figure 6.**
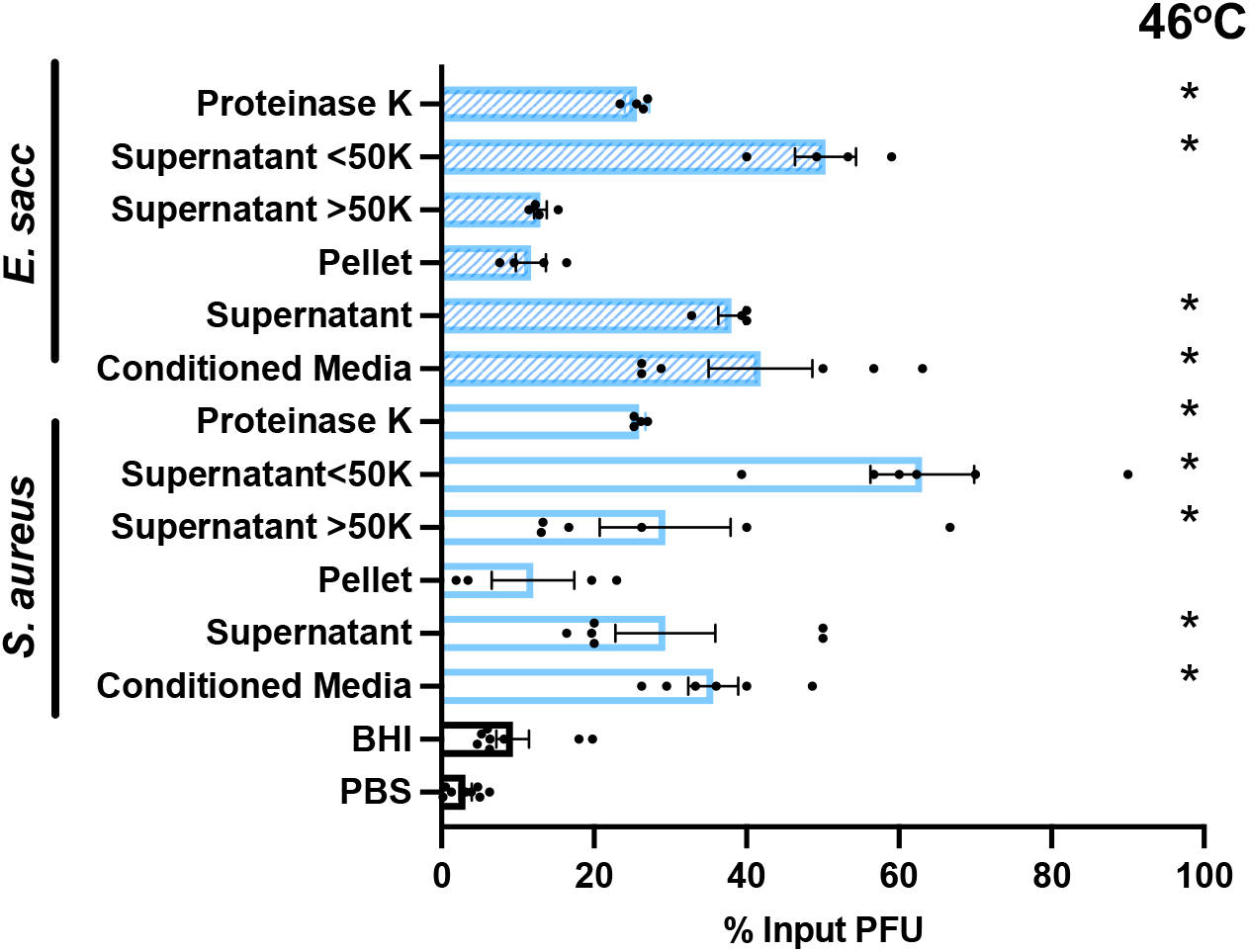
Effects of conditioned media treatments on stabilization of MNV. Thermal stability assays were performed by incubating 1 x 10^6^ PFU MNV with PBS, BHI, and conditioned medium from *S. aureus* and *E. saccharolyticus* that was filtered with a 0.2 μm filter and then separated with 50 kD spin column or subjected to ultracentrifugation at 150,000 x g for 3 h to generate supernatants and pellets. The samples were incubated with virus at 46 °C for 4 h. The amount of viable virus after each assay was determined by plaque assay and compared to a 4°C PBS control to calculate percent of input PFU that remained. Data are representative of 2 to 4 independent experiments *(n* = 4 to 8). Bars show mean and SEM. Statistical significance was determined by one-way ANOVA compared to PBS (*, *P* < 0.05).

Lastly, we found that the stabilizing effects of conditioned media on MNV did not increase infectivity in a plaque assay (Fig. 4). This may indicate that bacteria play more of a role in maintaining viability for a specific viral particle in a certain environment or length of time rather than making the particle more infectious. Other enteric viruses such as coxsackievirus B3 and poliovirus have a correlation between increased infectivity and stability (10, 13). This may indicate that the mechanism for MNV stabilization is distinct from other enteric viruses.

Overall this study illuminates the role that specific bacterial species and bacterial surface components play in MNV infection and uncovers MNV stabilization by bacterial conditioned media. Understanding the role of bacterial species and bacterial conditioned media in stabilizing MNV can provide insight into how MNV establishes an infection and how MNV may spread between hosts due to increased stability in the environment.

## Materials and Methods

### Cells and Viral Stocks

BV2 cells were grown in Dulbecco’s Modified Eagle Medium (DMEM) with 10% fetal bovine serum, 1% HEPES, and 1% penicillin-streptomycin. MNV-1.CW3 (MNV1) (5) was generated by transfecting HEK293T cells with an infectious clone plasmid followed by two rounds of amplification in BV2 cells to generate high titer viral stocks. Viral stocks were stored at −80°C.

To quantify virus, plaque assays were performed as previously described. Briefly, virus was diluted in phosphate-buffered saline supplemented with 100 μg/ml CaCl_2_ and 100 μg/ml MgCl_2_ (PBS+) and added to BV2 cells for 30 min at 37°C to facilitate attachment. Overlays containing 7.5% methylcellulose, MEM, 10% fetal bovine serum, 1% penicillin-streptomycin, and 1% HEPES were used and removed after 72 h.

Radiolabeled virus was generated by propagating BV2 cells in the presence of [^35^S]cysteine-methionine. BV2 cells plated in 15 cm plates were infected at a MOI of 0.05 for three hours before adding 0.36 mCi of [^35^S]cysteine-methionine (Perkin Elmer) for 45 h. The media and cells were freeze thawed 3 times and centrifuged to remove cell debris. Supernatants were centrifuged through a 30% sucrose cushion at 27,000 rpm for 3 h at 4°C in SW28 rotor. Viral pellets were resuspended in 350 uL of 10% N-lauryl sarcosine and left at room temperature for 2 hours. Virus was added to CsCl solution made in PBS and adjusted to a refractive index of 1.3665. CsCl gradients were formed by ultracentrifugation at 35,000 rpm for 40 h at 12°C in SW55 rotor. Individual fractions were collected from the top of the gradients followed by scintillation counting and performing plaque assay to determine virus-containing fractions. The purity of virus was confirmed by SDS-PAGE and phosphor imaging. Viral fractions were dialyzed against PBS at 4°C before storing at −80°C in glass tubes.

### Bacterial Strains and Bacterial Conditioned Media

Bacterial strains were from ATCC or from the cecum of mice as described previously (11). Note that the *E. cloacae* strain used here is a non-ATCC strain from a teaching lab and has unknown H antigen status (5, 11). Overnight cultures were inoculated from glycerol stocks in BHI media. The OD_600_ value was determined for each culture by spectrophotometer (Eppendorf BioPhotometer D30) to determine the CFU required for each experiment. The required volume of bacteria was pelleted and washed two times and resuspended in PBS+. Bacteria were UV inactivated prior to use in assays by exposing the bacteria to UV light for 30 minutes. UV inactivation conditions were confirmed by plating on BHI agar.

Bacterial conditioned media was generated by growing overnight cultures of each bacterial strain before different types of processing for each conditioned media experiment. For Figure 5A, bacteria were pelleted by centrifugation and the supernatant is referred to as “spent media”. The spent media was then either filtered using a 0.2 micron filter or boiled in a 95°C heat block for 30 minutes. For Figure 5B, bacteria were pelleted and the supernatant was then filtered with a 0.2 micron filter. The filtered media is referred to as “conditioned media”. For Figure 6, the conditioned media was spun at ~150,000 x g for 3 h at 4°C in an ultracentrifuge. The supernatant was collected and the pellet was resuspended in PBS+. The supernatant was then separated using a 50kD spin column (Millipore Sigma) and both the >50kD and <50kD fractions were collected. The conditioned media was also treated with 0.1 mg/ml proteinase K (Invitrogen Ambion) for 18 h at 37°C (24). The enzyme was inactivated by boiling the sample for 30 min.

### Viral Stability Assays

To determine whether bacteria impacted the stability of MNV, 1 x 10^6^ PFU of MNV was mixed with either PBS+, 1 x 10^9^ CFU bacteria, 1 mg/mL of compounds/molecules, or 200 uL bacterial conditioned medium and incubated at 42°C for 6 h or 46°C for 4 h. A control sample of virus in PBS+ was placed at 4°C for the duration of the experiment. After incubation plaque assays were performed on both the heat exposed samples and the 4°C control sample using BV2 cells to determine the amount of viable virus before and after heat treatment. The percent of input PFU remaining after heat exposure was calculated by dividing the titer of each sample by the control sample.

### Binding Assays

The bacterial binding assay was performed as previously described for poliovirus. Approximately 3,500 cpm of ^35^S-radiolabeled virus was mixed with PBS or 1 × 10^9^ CFU of bacteria and incubated at 37°C for 1 h. Binding reactions were done in the presence of 0.1% BSA to prevent nonspecific binding. After incubation, bacteria were pelleted and washed with PBS+ to remove any unbound virus. CPM counts for both input and bacteria were obtained by scintillation counting to determine the amount of virus that was bound to bacterial cells.

### Infectivity Assays

To determine the effect of bacterial conditioned media on the infectivity of MNV, 1 x 10^6^ PFU was incubated with either PBS or conditioned media for 1 h at 37°C. Ten-fold dilutions of the pre-incubated MNV media were plated as in a standard plaque assay. The plaque assay plates were incubated for either 1 minute, 5 minutes, or 15 minutes, washed, and overlay was added. The titer of the virus incubated with PBS or the conditioned media was compared for each time point.

## Data Analysis

Statistical analyses were performed using GraphPad Prism software. All one-way ANOVAs were performed with Dunnett’s multiple-comparison *post hoc* test.

## Acknowledgements

We thank Andrea Erickson for critical review of the manuscript. Work in J.K.P.’s lab is funded through NIH NIAID grant R01 AI74668, a Burroughs Wellcome Fund Investigators in the Pathogenesis of Infectious Diseases Award, and a Faculty Scholar grant from the Howard Hughes Medical Institute. M.R.B. was supported in part by the NIH Molecular Microbiology Training Grant T32 AI007520.

